# Identification of Tumor-Reactive B Cells and Systemic IgG in Breast Cancer Based on Clonal Frequency in the Sentinel Lymph Node

**DOI:** 10.1101/212308

**Authors:** Jonathan R. McDaniel, Stephanie C. Pero, William N. Voss, Girja S. Shukla, Yujing Sun, Sebastian Schaetzle, Chang-Han Lee, Andrew P. Horton, Seth Harlow, Jimmy Gollihar, Jared W. Ellefson, Christopher C. Krag, Yuri Tanno, Nikoletta Sidiropoulos, George Georgiou, Gregory C. Ippolito, David N. Krag

**Affiliations:** Department of Surgery, Vermont Cancer Center, University of Vermont Larner College of Medicine, Burlington VT, USA; Department of Chemical Engineering, The University of Texas at Austin, Austin TX, USA; Department of Pathology and Laboratory Medicine, University of Vermont Larner College of Medicine, Burlington VT, USA; Department of Molecular Biosciences, The University of Texas at Austin, Austin TX, USA

**Keywords:** breast cancer, antibody, sentinel node, next-generation sequencing, repertoire

## Abstract

A better understanding of antitumor immune responses is key to advancing the field of cancer immunotherapy. Endogenous immunity in cancer patients, such as circulating anticancer antibodies or tumor-reactive B cells, has been historically yet incompletely described. Here, we demonstrate that tumor-draining (sentinel) lymph node (SN) is a rich source for tumor-reactive B cells that give rise to systemic IgG anticancer antibodies circulating in the bloodstream of breast cancer patients. Using a synergistic combination of high-throughput B-cell sequencing and quantitative immunoproteomics, we describe the prospective identification of tumor-reactive SN B cells (based on clonal frequency) and also demonstrate an unequivocal link between affinity-matured expanded B-cell clones in the SN and antitumor IgG in the blood. This technology could facilitate the discovery of antitumor antibody therapeutics and conceivably identify novel tumor antigens. Lastly, these findings highlight the unique and specialized niche the SN can fill in the advancement of cancer immunotherapy.

**SIGNIFICANCE:** Using high-throughput molecular cloning and antibody proteomics to study coordinated antitumor immunity in breast cancer patients, we simultaneously demonstrate that the sentinel lymph node is a localized source of expanded antitumor B cells undergoing affinity maturation and that their secreted antibodies are abundant as systemic IgG circulating in blood.

## INTRODUCTION

The recent success with cancer immunotherapy has sparked a renewal of interest in harnessing the immune system for tumor eradication and for the identification of novel tumor antigens in cancer patients. Underlying most approaches to cancer immunotherapy is a pivotal reliance on a complex and enigmatic repertoire of tumor-reactive adaptive immune cells that have been exposed to tumor antigens. The advent of next-generation sequencing and advanced proteomics technologies may allow a comprehensive characterization of the localized repertoire of tumor-reactive lymphocytes found in the tumor or lymph node as well as identification of systemic B-cell-derived circulating antibodies (IgG and IgA). This represents a unique opportunity to discover patient-specific, tumor-reactive cell repertoires, to study their temporal dynamics and antitumor function, and to potentially reveal the tumor antigens they target *in vivo*. At present, however, there is limited knowledge of tumor-reactive immune repertoires in cancer patients, and it is generally not known to what extent an adaptive immune response is activated in any individual cancer patient when treated with cancer immunotherapy such as immune checkpoint blockade. Given the complexity of the relationship between cancer and the immune system, a better understanding of the parameters of human adaptive immunity and its potential role in the eradication of solid tumors can ultimately be expected to lead to the design of better, personalized therapies.

Preclinical and clinical evidence suggests that breast cancer is under immune surveillance, that the majority of patients with breast cancer have some evidence of adaptive antitumor immunity (1), and that current regimens are most effective if they elicit a robust tumor-targeting immune response (2). The prognosis of breast cancer patients can be significantly correlated with the density and composition of the immune infiltrate at diagnosis, especially in the triple negative breast cancer (TNBC) subtype. The most recent clinical trial data for TNBC—long considered to be an improbable candidate for immunotherapy—confirms that TNBC is immunogenic and responsive to PD-1 checkpoint immune modulation (3,4). The anatomic distribution of lymph nodes in breast cancer patients offers a unique glimpse into the dynamics between the tumor and the adaptive immune response as breast cancer lymph, rich in tumor antigens, typically drains directly to a small cluster of interconnected lymph nodes aptly termed tumor-draining lymph nodes or sentinel nodes (SNs) (5,6). Recently, there has been significant interest in mining the B cell receptor sequences from SNs, as the presence of tumor antigens is likely to drive affinity maturation of anticancer antibodies (7,8).

We have developed a high-throughput platform for the rapid identification of endogenous antibodies arising during a cancer patient's adaptive immune response to tumor (Figure 1). This technology is a synergistic combination of IgG protein mass spectrometry (Ig-Seq) and a DNA sequencing method that preserves the natural pairing of heavy (VH) and light (VL) chain variable regions isolated from hundreds of thousands of B cells in a single experiment (VH:VL BCR-Seq) (9–11). This combined approach allows for (i) the facile discovery and characterization of antigen-specific recombinant monoclonal antibodies (mAbs), and (ii) the identification and semi-quantitation of these IgG or IgA antibodies in blood.

**Figure 1.**
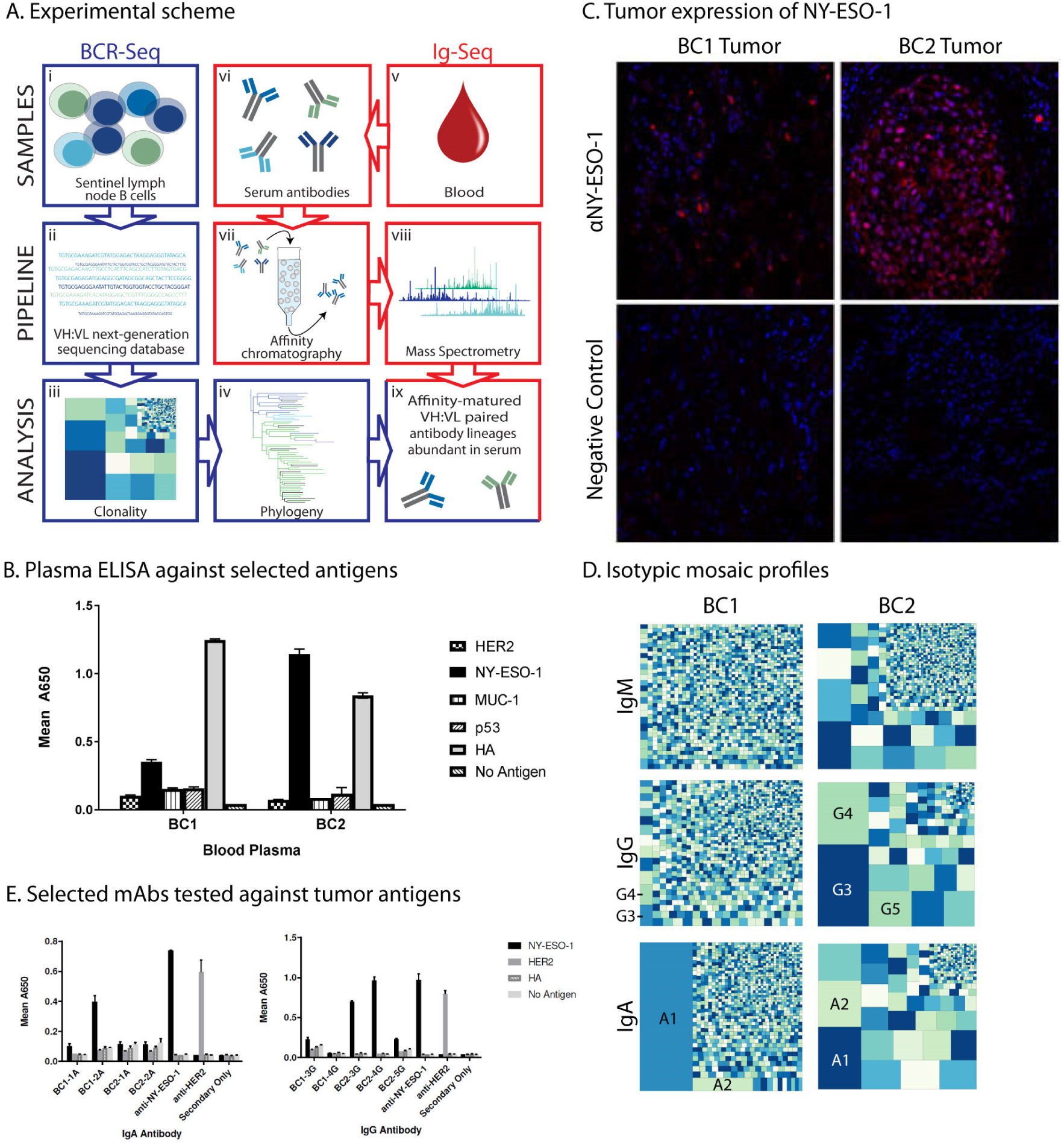
(A) Identification of antibodies from the SN B cells and blood of breast cancer patients. **(i)** BCR-Seq utilizes patient-specific B cells to **(ii)** build a patient-specific next-generation VH:VL paired sequencing database. The database is then used to **(iii)** identify highly expanded lineages by mosaic profiling and **(iv)** construct phylogenetic trees to describe clonal relationships. **(v)** The Ig-Seq pipeline isolates **(vi)** total IgG from plasma and **(vii)** uses affinity chromatography to enrich the sample for antibodies against a specific antigen. **(viii)** The IgG molecules are cleaved into peptides, run on LC-MS/MS, and **(ix)** identified by mapping the MS spectra to the sequencing database. **(B)** An ELISA was used to identify NY-ESO-1 sero-positive patients by screening blood plasma. BC1 and BC2 were selected for further characterization due to the presence of anti-NY-ESO-1 antibodies. The antigen panel included: HER2, NY-ESO-1, MUC-1, p53, and Hemagglutinin Flu b (HA) protein as a positive control. Values represent mean ± SD (n=2). **(C)** NY-ESO-1 expression was detected in the tumor tissue of breast cancer patients BC1 and BC2. Representative fluorescent images of tumor tissue stained with anti-NY-ESO-1 followed by Alexa fluor 555 conjugated goat anti-mouse-IgG (H+L) secondary (top images) or secondary antibody only (bottom images). **(D)** Mosaic profiles of expanded B cell clones in the SN. The size of each square is relative to the frequency of NGS reads assigned to each lineage following down-sampling to normalize the number of reads across patients and isotypes. Expressed antibody lineages are labeled. **(E)** Selective human antibodies derived from the SN B cells binding to NY-ESO-1. ELISA demonstrating strong positive IgA (BC1-2A) and IgG (BC2-3G and BC2-4G) antibodies binding to NY-ESO-1 from patient BC1 and BC2 and not to HER2 or hemagglutinin Flu B (HA) protein. Values represent mean ± SD (n=2).

As a model tumor antigen for exploring the utility of the BCR-Seq/Ig-Seq platform, we chose NY-ESO-1. This tumor antigen was first identified using tumor mRNA and autologous patient blood (12), is one of the most immunogenic human cancer antigens known to date (13), and is expressed in up to 25% of breast tumors (14). NY-ESO-1 has expression in primary breast carcinoma correlated with plasma cell infiltration (15), and pre-existing serological immunity to NY-ESO-1 in metastatic melanoma and non-small-cell lung carcinoma has been shown to both correlate with improved clinical benefit following anti-CTLA-4 checkpoint immunotherapy and with CD8+ T cell responses (16).

Given that SN is exposed to tumor-associated antigens like NY-ESO-1, and therefore presumably enriched for tumor-reactive B-cell clones, we tested the hypothesis that the most highly expanded B-cell clones will be tumor-reactive. We also tested the additional hypothesis that SN B-cell clonal expansions might be detectable in peripheral blood as circulating immunoglobulin (IgG). Using the combined VH:VL BCR-Seq and Ig-Seq technologies, we present data establishing how chronic exposure to tumor antigens drives quantitative clonal expansion and affinity maturation of tumor-specific B cells localized within SNs; that these tumor-specific B cell clones may be identified prospectively using high-throughput sequencing; moreover, we establish that these SN B cells give rise to plasma cells and systemic, circulating antitumor IgG.

## RESULTS

### Patient selection

Newly diagnosed patients with breast cancer were evaluated for elevated antibody titers to a small panel of tumor-related antigens. At the time of surgery we collected blood, a portion of the SN, and a portion of the primary breast tumor. The blood plasma was measured for elevated antibody titers to HER2, MUC-1, p53, and NY-ESO-1. Hemagglutinin Flu B (HA) protein antigen was included as a positive control. We identified two breast cancer patients, BC-1 and BC-2, with elevated blood plasma titers to NY-ESO-1 antigen (Figure 1B). Both patients were diagnosed with invasive ductal carcinoma and their tumors were greater than 1 cm in size. The BC1 patient tumor was ER+/PR+/HER2- while the BC2 patient tumor was TNBC ER^-^/PR^-^/HER2^-^. Primary tumor tissue in patients BC1 and BC2 was confirmed to have positive NY-ESO-1 expression by immunofluorescence staining (Figure 1C).

### Interrogation of SN B cells

To test the hypothesis that highly expanded B cell clones are tumor-reactive, we isolated approximately 2-3 million mononuclear cells from the SN. These cells were subjected to paired BCR-Seq (17,18), a process by which the VH and VL antibody transcripts from a single B-cell antibody receptor (BCR) are reverse transcribed and physically linked (VH:VL) using emulsion-based overlap-extension RT-PCR. The RT-PCR reaction integrates reverse transcription xenopolymerase (RTX), an enzyme that exhibits both reverse transcriptase and polymerase activity (19), to enhance the fidelity and efficiency of the reaction. The VH:VL library was deep sequenced using paired-end Illumina MiSeq, bioinformatically filtered to remove low-confidence sequences, and the annotated antibody sequences were separated by isotype.

We then grouped the VH:VL antibody sequences into antibody lineages by applying a 90% similarity threshold on the CDR-H3 region, and randomly sampled 20,000 reads from each isotype to form representative mosaic plots (Figure 1D), where the area of each square is proportional to the number of sequencing reads belonging to each lineage.

In total, we identified a total of 17,855 lineages from the B cells isolated from BC1 SN (474 thousand productive VH:VL sequences; 1.7 million raw reads), and 8,678 lineages from the B cells isolated from BC2 SN (512 thousand productive VH:VL sequences; 1.7 million raw reads). Figure 1D illustrates vast differences in the degree of clonal expansion between each of the patients and isotypes. We observed that the class switched IgG and IgA isotypes displayed fewer lineages than the respective IgM compartments. Following 100 repetitions of the random sampling and lineage enumeration used for Figure 1D, the number of IgG and IgA lineages relative to IgM averaged 77% and 83% for BC1, respectively, and 27% and 15% for BC2. This polarization, which was most pronounced in BC2, suggests that the class-switched B cells from the SN are less diverse because they are experiencing antigen-driven clonal expansion. While this visualization strategy provides no information on the identity of the antigens driving the immunological response, we hypothesized that it may identify anticancer antibody lineages that experienced affinity selection in the SN germinal centers in response to chronic stimulation by tumor antigens.

### Phylogenetic analysis of resident B cells in the SNs

To identify the most promising antibody candidates for characterization, we mapped each SN sequencing database to a frequency-based mosaic plot as a first-pass filter to identify the clonally expanded and class-switched antibodies (Figure 1D). We first decided to examine only the most expanded clonal lineages in the class-switched compartments: from BC1, we selected 1A, 2A, 3G and 4G and from BC2 we chose 1A, 2A, 3G, 4G and 5G. Full-length heavy-chain sequences (VH genes) were constructed into maximum likelihood phylogenetic trees partitioned on the variable region IGHV gene segment. Clonal sub-lineages were then defined by clusters of VH genes containing identical CDR-H3 amino acid sequences.

In a remarkable display of lineage diversification, the BC2-3G phylogram diverged into 4 sub-lineages (differentiated by color), each defined by a related, but unique, CDR-H3 amino acid sequence originating from the joining of the VH6-1, DH6-13, and JH4 gene segments (Figure 2A). Furthermore, the paired sequencing data in Figure 2B suggests that each CDR-H3 sub-lineage preferred a specific CDR-L3 variant originating from a VK3-11 and JK4 light chain phylogeny. The observation that specific mutations appear linked across both VH and VL genes is highly suggestive of clonal expansion and somatic hypermutation consistent with antigen-driven affinity maturation in germinal centers (Figure 2C, D). BC2-4G and BC2-5G clonal lineages also displayed this behavior, each with 2 major sub-lineages that utilized a single VH and VL gene combination (SI Figures 1-2).

**Figure 2.**
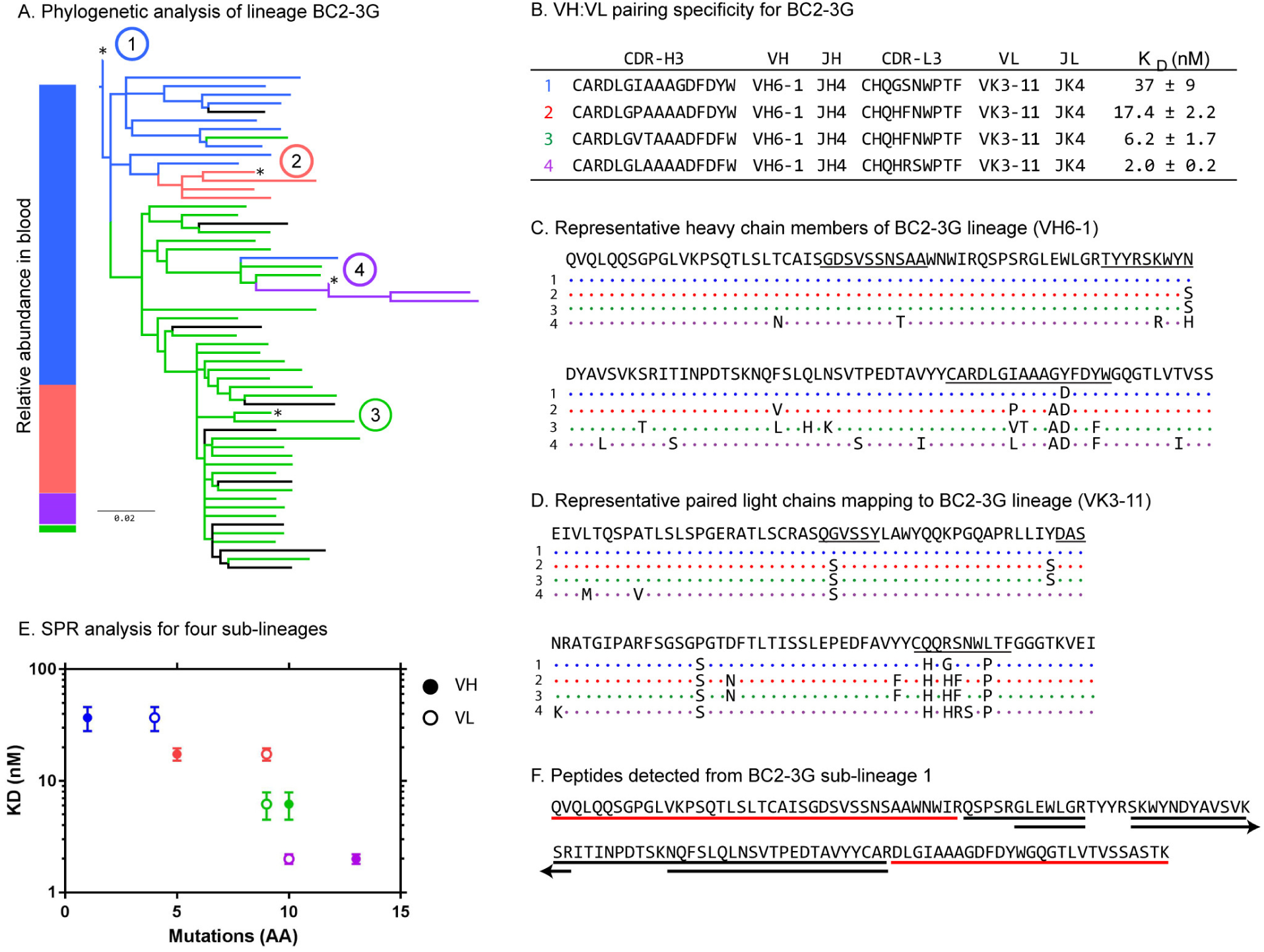
Antigen-driven affinity maturation of a B-cell lineage. **(A)** Maximum likelihood phylogram of BC2-3G. Sub-lineages are defined by different CDR-H3 amino acid sequences, and are differentiated by color. Black nodes represent CDR-H3 sequences that do not exactly match one of the primary sub-lineage CDR-H3s. Side bar represents the relative abundancies of each color-coded sub-lineage detected in blood. **(B)** Table indicating the pairing specificity of CDR-H3 to CDR-L3 across dominant lineage members and affinity for NY-ESO-1. Representative **(C)** VH and **(D)** VL sequences for each sub-lineage denoted by an asterisk in (A). CDR1, CDR2, and CDR3 regions are underlined. **(E)** Relationship between the number of somatic mutations in the VH and VL genes and K_D_ as determined by SPR for lineage members 1 (blue), 2 (red), 3 (green), and 4 (purple). K_D_ values are shown in Table B. **(F)** Underlined sequences represent peptides detected by LC-MS/MS that map to sub-lineage 1. Red indicates that the peptide was unique to this lineage, black indicates that the peptides may have derived from several lineages. Only the CDR-H3 peptide was used to quantify relative abundancies between lineages.

Having demonstrated the occurrence of highly expanded B-cell clones (VH:VL pairs) in the SN, we asked whether such clones could encode antibodies reactive with breast cancer tumor. To this end, we recombinantly expressed representative VH:VL pairs as full-length antibodies in HEK293 cells and screened via ELISA (Figure 1E) against recombinant NY-ESO-1 as both patients BC1 and BC2 displayed elevated plasma titers to this antigen. Additional sub-lineages from each of the positive hits were then expressed and examined via surface plasmon resonance (SPR). Monoclonal antibodies from BC2-3G and BC2-4G lineages displayed nM affinity for binding to NY-ESO-1 as determined by SPR analysis (SI Figure 3A-E), with the apparent affinities varying greatly between the lineages (SI Figure 3F).

The highest affinity monoclonal antibody screened by ELISA was a member of the BC2-3G phylogeny; surprisingly, the entire IGHV gene segment matched germline VH6-1, and even the CDR-H1 and CDR-H2 were unmutated (labeled as (1) in Figure 2A). To investigate whether antigen-driven somatic hypermutation was able to improve the affinity of this germline sequence (blue), we expressed sub-lineages 2 (red), 3 (green) and 4 (purple), and examined all four mAbs by SPR against commercial NY-ESO-1. As further evidence of clonality among the sub-lineages of BC2-3G, all four recognized NY-ESO-1 with varying affinities. Supporting a narrative of active, affinity-driven selection, we observed a strong correlation between the number of mutations in the variable region of the VH and VL genes and the affinity of the antibodies (as measured by K_D_), shown in Figure 2E.

### Identification of SN mAbs as circulating IgG in peripheral blood

Finally, we wanted to determine if the most expanded IgG transcripts isolated from the SN were represented by antibodies present in the blood immunoglobulin (Ig). To elucidate the identity of circulating antibodies from patient BC2, we performed bottom-up proteomics on plasma IgG using a process termed Ig-Seq (20–22). Following plasma IgG enrichment and proteolytic cleavage of the hinge region, NY-ESO-1-specific F(ab’)2 was isolated with affinity chromatography, digested with trypsin and subjected to LC-MS/MS. The resulting spectra were identified using the donor specific B-cell receptor sequence database generated from the SN (Figure 1A). In this experiment, the abundance of each lineage was measured by the intensity of the CDR-H3-J region peptide. Using this approach, we confirmed the presence of BC2-3G and BC2-4G lineages in the plasma. The relative abundances of the plasma response to NY-ESO-1 indicated the presence of all 4 distinct sub-lineages of the BC2-3G lineage (shown in Figure 2A) and both sub-lineages of BC2-4G. Figure 2F illustrates the observed peptide coverage for BC2-3G lineage 1. Whereas the CDR-H3 peptide is unique to the BC2-3G lineage (indicated by a red underline), many other peptides were observed across multiple lineages due to their low mutation rate (indicated by black underlines). However, it should be noted that due to current limitations in mass spectrometry protein identification, only lineages present in the SN at the time of resection could be identified. Using an in-house spectral classifier, we estimate that this analysis identified ~33% of the CDR-H3-J region peptide spectra observed through LC-MS/MS.

## DISCUSSION

Using NY-ESO-1 tumor antigen as a paradigm, we have comprehensively assayed the humoral immune response to breast cancer by analyzing the B-cell repertoire expanded in the SN and the antigen-specific Ig repertoire in blood plasma using next-generation sequencing and proteomics (Figure 1A). We determined the molecular composition of polyclonal immunoglobulins binding NY-ESO-1 and identified B-cell clonal lineages in the SN responsible for the production of these anticancer antibodies. Circulating anticancer IgG antibodies were then quantitated (at monoclonal resolution) and linked conclusively to radically expanded, affinity matured clones residing among total SN B cells. The expansion of SN B cell clones could be represented graphically using mosaic plots, which enabled the prospective identification of tumor-reactive B cells based solely on clonal frequency; these were subsequently confirmed to be specific for NY-ESO-1 when expressed and tested as high-affinity recombinant monoclonal antibodies (mAbs).

While B cells from SNs have been analyzed in previous studies using low-throughput techniques (8,23,24), here we used next-generation sequencing to provide a comprehensive high-throughput survey of SN B cells; moreover, we obtained natively paired VH and VL sequences at high throughput from SN B cells and validated their tumor specificity and high-affinity binding. This distinction is vital as it provides a depth of coverage that allows us to observe the clonality and phylogeny of the resident B cells, and to infer tumor specificity based upon the extent of clonal expansion. Importantly, because our technology can identify antitumor IgG circulating in plasma that is clonally related to expanded SN B-cell lineages, we can infer which SN B cells give rise to antibody-secreting plasma cells. Our results suggest there is systemic coordination of antitumor immunity and immune activation between a tumor-reactive lymph node and the periphery, in agreement with a recent precedent established for mouse models of tumor immunity (25). We conclude that chronic exposure to tumor antigens drives clonal expansion and affinity maturation of B cells localized within SNs, which give rise to plasma cells and systemic antitumor IgG. The ability to identify these trained B cell receptors could potentially lead to prognostic indicators and therapeutic antibodies against cancer.

Because high-throughput analysis of antibody transcripts (native VH:VL pairs) using paired sequencing technology (VH:VL BCR-Seq) (17,18) enables frequency-based and phylogenetic analysis of full-length antibodies, this approach potentially obviates the need for constructing combinatorial display libraries that may bias the frequency and identity of the antibody lineages present in the SN. We identified prospective B-cell lineages to be clonally expanded as evidenced by the sheer number of cognate VH and VL transcripts outputted by the next-generation sequencing reaction. Furthermore, each of the top B-cell lineages examined could be categorized into multiple VH sub-lineages differentiated solely by sequence variability in a key peptide interval, the CDR-H3 region, long regarded as an identifying “fingerprint” of B-cell progenitors and their progeny; CDR-H3 lies at the center of the antibody binding site, is known to be a mutation hotspot for affinity maturation, and is a primary determinant of antibody specificity (26,27). Third, we observed a higher frequency of mutations focused on all of the CDR regions when compared to the framework regions, and also a high rate of mutations shared between the VH and VL sub-lineages (Figure 2C-D), indicating that these mutations could not result from randomly distributed PCR or sequencing errors. It should be noted that our analysis did not exclude any B cell phenotype; the process was performed on the mixture of total B cells, whether antibody-secreting plasmablasts, memory B cells, or naïve B cells present in the SN. PCR bias did not influence the number of reads per lineage, for we observed a strong correlation between the frequency of V-gene usage in the top 1000 and bottom 1000 lineages ranked by read count (Spearman ρ=0.95).

In addition to these sequence and phylogenetic analyses, we also demonstrated empirically that the B cells in an expanded lineage had experienced affinity-based maturation and selection. We expressed four mAbs representative of each VH sub-lineage and characterized their affinity to the tumor antigen NY-ESO-1. We observed a clear stepwise correlation between affinity for the antigen and the number of VH and VL somatic mutations. Thus, screening for expanded and branched B cell phylogenies isolated from a SN may prove a useful approach to discover affinity-matured antibodies even with no *a priori* knowledge of antigen-specificity. While the strategy presented in this paper highlights antibody lineages that recognize the NY-ESO-1 tumor antigen, it follows that a target-agnostic approach of mining the SN for autoantibodies could be applied to other known tumor antigens as well as those yet to be discovered. Although SNs are not present in every form of cancer, we believe that this technique can be applied to a variety of relevant B cell populations, including those isolated from ectopic germinal centers and tumor tissue. Slight modifications to the VH:VL BCR-Seq technology have also allowed us to analyze the natively paired T cell receptor (TCR)β:TCRα repertoire at high throughput (manuscript in preparation), suggesting that we may soon be able to determine the cognate BCR and TCR antitumor repertoire from SNs. Similar to our observations with B cells, we anticipate discovery of NY-ESO-1-specific T cells identifiable by clonal expansion in the SN (28,29). This may prove a significant step forward in the development of patient-specific combination adoptive cell therapies. NY-ESO-1 is an attractive candidate for immunotherapy in TNBC (30) and early-phase clinical trials of NY-ESO-1 TCR immunotherapy have shown remarkable 55%–80% objective clinical response rates for melanoma, synovial sarcoma, and multiple myeloma (31,32).

Lastly, the ability of Ig-Seq proteomics to identify and quantitate antibody proteins circulating in patient blood enables a molecular-level analysis of the monoclonal antibodies that comprise a tumor-reactive immunoglobulin (Ig) repertoire and opens new vistas for cancer research. Whereas our approach was sufficient to demonstrate that highly expanded B cell lineages are clonally related to antitumor IgG circulating in the blood, a complete accounting of the tumor-specific serological repertoire would likely require a comprehensive database assembled from multiple tissues in addition to the SN—including primary tumor, bone marrow, and peripheral blood, which will be the focus of future studies. As a diagnostics tool, we envision that Ig-Seq humoral monitoring may be predictive of clinical outcomes, such as the prediction of partial versus complete responses (versus stable disease) for patients treated with immune checkpoint blockade. As a discovery platform, Ig-Seq could facilitate the discovery of antitumor antibody therapeutics, innately endowed with optimized drug-like properties. The platform could also conceivably identify novel tumor antigens, including tumor mutations, using the patient-specific affinity-selected antitumor antibodies as bait; this approach holds promise to address the current deficit of targets delimiting the field of immuno-oncology.

## METHODS

### Tissue Acquisition and Processing

The collection of tumor tissue and blood was performed under protocols approved by the Institutional Review Board at the University of Vermont. After informed consent was obtained, SNs were resected using radiotracer guidance to accurately locate the SN receiving direct drainage from the breast cancer (33,34). The freshly harvested SNs were minced in cold RPMI-10% FBS media between 40 µm nylon mesh and pressed with a rubber plunger to isolate mononuclear cells. The number and viability of cells were determined using trypan blue exclusion, and the cells were cryopreserved in 90% fetal bovine serum (Sigma)/10% dimethyl sulfoxide. The blood was collected in Monoject™ EDTA coated blood collection tubes (Covidien) at time of surgery and then transferred to a centrifuge tube. The plasma layer was collected by centrifugation (1000 g for 10 min). The upper plasma layer was stored in aliquots at -20 °C.

### Binding Analysis of Plasma and Purified Antibodies by ELISA

Flat-bottom MaxiSorp 96-well plates (Nunc) were coated overnight at 4°C with antigens (5 µg/mL) in phosphate buffered saline (PBS), or PBS alone. Human antigens include: extracellular domain of ErbB2 (HER2; Creative Biomolecules Inc.), MUC-1 (partial ORF 315-420 aa; Abnova), Hemagglutinin (HA; FluB/Florida/4/2006; eEnzyme), NY-ESO-1 (Thermo Fisher or RayBiotech Inc.), and p53 (Sigma-Aldrich). Wells were thoroughly washed with Tris-buffered saline containing 0.1% Tween-20 (TBST-0.1%) and nonspecific binding sites were blocked for 2 hours with 1% bovine serum albumin (BSA; Sigma-Aldrich). Diluted plasma (1:2000 in PBS) or purified human antibodies was incubated in antigen coated wells for 2 h at room temperature. Human anti-HER2 (trastuzumab; Genentech Inc.) and mouse anti-NY-ESO-1 monoclonal antibody (E978; Thermo Fisher) were used as positive controls. After washing the plates, antigen-reactive antibodies were detected with mouse anti-human IgG (Fc)-horseradish peroxidase (1:5000 dilution; Southern Biotech) or goat anti-mouse IgG (H+L)-horseradish peroxidase conjugate in Casein-TBS Blocker (Thermo Fisher) with gentle shaking for 1 hour. After washing 5 times with TBST-0.1% and once with TBS, the bound antibody was detected with 3,3’,5,5’-tetramethylbenzidine soluble substrate (TMB; Millipore)by monitoring the formation of blue-colored product at 650 nm for 15 min using a Synergy HT plate reader (BioTek Instruments, Inc.).

### Immunofluorescence microscopy of tumor NY-ESO-1

Frozen sections (6 µm thick) mounted onto uncoated glass slides were fixed in 4% paraformaldehyde (Electron Microscopy Sciences) in PBS, rinsed in PBS, and blocked in 10% normal goat serum (Jackson Immuno Research) diluted with PBS containing 5.0% BSA and 0.1% Triton X-100. All washes were performed with PBS containing 5% BSA. Tumor NY-ESO-1 was detected with 2.5 µg/mL anti-NY-ESO-1 monoclonal antibody (Invitrogen), followed by 4 µg/mL Alexa fluor 555-conjugated goat anti-mouse-IgG (H+L) (Invitrogen). Representative images were obtained on the 510 META confocal scanning laser microscope (Zeiss).

### Sequencing the paired VH:VL repertoire

The VH and VL variable regions were determined on a single cell basis using a custom-designed axisymmetric flow focusing device as previously described (17,18), with few modifications. B cells were purified from the isolated SN cells using the human B cell enrichment kit (StemCell Technologies) according to the manufacturer’s instructions. Briefly, the flow focusing device was used to coemulsify single cells with lysis buffer (100 mM Tris pH 7.5, 500 mM LiCl, 10 mM EDTA, 1% lithium dodecyl sulfate, and 5 mM DTT) and oligo d(T)_25_ magnetic beads (New England Biolabs). The magnetic beads were washed, resuspended in a customized high-yield RT-PCR solution (19) (SI Table 1), emulsified, and subjected to overlap-extension RT-PCR under the following conditions: 30 min at 68°C followed by 2 min at 94°C; 25 cycles of 94°C for 30 s, 60°C for 30 s, 68°C for 2 min; 68°C for 7 min; held at 4°C. The RT-PCR solution contained a multiplex set of VH and VL primers (SI Table 2) designed to physically stitch the two antibody chains into a single amplicon. The resultant VH:VL DNA amplicon was isolated from the emulsions, amplified using nested PCR, and then split into three libraries for paired-end sequencing by Illumina: the full length VH and VL amplicon to maintain the CDR-H3:CDR-L3 paired information, and the separate VH and VL libraries to provide phylogenetic data on the full length variable region.

### Bioinformatic analysis

#### VH:VL repertoire analysis

Raw 2x300 MiSeq reads from the paired VH:VL library were trimmed based on sequence quality (35) and submitted to MiXCR for gene annotation (36). Productive VH and VL reads were split by isotype and paired using a custom python script. Sequences with ≥2 reads were grouped into lineages by clustering the CDR-H3 region on 90% nucleotide identity. The number of Illumina reads for each lineage was transformed into a frequency to generate mosaic plots for each isotype (37).

#### Phylogenetic analysis

Raw MiSeq reads were stitched into full length variable regions using PEAR (38), quality filtered, and annotated by MiXCR (36). Lineages of interest were identified by 90% CDR-H3 nucleotide identity, and their members were clustered on the full length nucleotide sequence at 97% identity to reduce PCR and sequencing error. The sequences were aligned by MAFFT (39) and organized into maximum likelihood phylogenetic trees based on the variable region using RAxML (40). Sub-lineages were defined on the basis of the CDR-H3 amino acid sequence.

#### Ig-Seq sample preparation and mass spectrometry

Total IgG (7.5 mg) was isolated from 1 mL plasma using Protein G Plus agarose (Thermo Fisher Scientific) affinity chromatography and cleaved into F(ab’)2 fragments using IgeS (FabRICATOR; Genovis). NY-ESO-1 specific F(ab’)2 was isolated by affinity chromatography using recombinant NY-ESO-1 coupled to NHS-activated agarose resin (Thermo Fisher Scientific). F(ab’)2 was applied to the column in gravity mode with the flow-through collected. Following elution with 100 mM glycine-HCl pH 2.7 and neutralization, the protein-containing fractions were pooled and prepared for LC-MS/MS as described previously (9). Briefly, the antigen-specific elution and the non-specific flow-through were separately concentrated, denatured in 50% (v/v) TFE and 10 mM PBS, reduced with 2.5 mM DTT, and incubated at 55˚C for 45 min. The samples were then alkylated with 15 mM iodoacetamide (Sigma) for 30 min at RT in the dark. Each sample was diluted 10x with 40 mM Tris-HCl pH 8.0 and digested with trypsin (1:50 trypsin/protein) for 4 hrs at 37˚C. Formic acid (1% v/v) was used to quench the digestion. The peptides were then concentrated under vacuum, resuspended in 5% acetonitrile and 0.1% formic acid, and washed on C18-SpinTips (Thermo Fisher Scientific) according to the manufacturer’s instructions. Finally, the peptides were analyzed by liquid chromatography tandem mass spectrometry (LC-MS/MS) using a Dionex UltiMate 3000 RSLCnano reverse phase chromatography system (Dionex Acclaim PepMapRSLC C18 column; Thermo Scientific) coupled to an Orbitrap Velos Pro mass spectrometer (Thermo Scientific). Parent ion MS1 scans were collected in the Orbitrap at 60,000 resolution. Ions with >+1 charge were fragmented by collision-induced dissociation (NCE 35), with a maximum of 20 MS2 spectra collected per MS1. Ions selected twice in a 30-second window were dynamically excluded for 45 seconds. Both the elution and the flow through were run three times.

#### MS/MS data analysis

A patient-specific antibody database was constructed by appending the full length IgG VH sequences to a database of background proteins comprising patient-specific VL sequences, Ensembl human protein-coding sequences, and common contaminants (maxquant.org). The spectra were then searched against this database using SEQUEST (Proteome Discoverer 1.4, Thermo Scientific) as described (41). High confidence peptide spectrum matches (PSMs) were filtered with Percolator (Proteome Discoverer1.4) at a false discovery rate of <1%, and only peptides with an average mass deviation <1.5 ppm were retained for downstream analysis. The full-length VH IgG sequences were organized into protein groups using a single-linkage hierarchical clustering algorithm requiring ≥90% amino acid identity in the CDR-H3 as measured by edit distance (41); peptides mapping to only a single protein group were considered informative. The relative abundances of the corresponding protein groups were calculated as a sum of the extracted-ion chromatogram (XIC) peak area for informative CDR-H3-J region peptides.

We also evaluated the MS/MS spectra using a custom machine learning system trained to recognize CDR-H3-J region peptide spectra. Built on a random forest classifier, this was trained in-house against a library of ~100,000 positive (CDR-H3-J) and negative (not CDR-H3-J) spectra collected in previous repertoire studies. Given an unknown spectrum, it predicts whether that spectrum derived from a CDR-H3-J peptide. It performs independently of any sequence database and does not predict the peptide sequence.

#### Antibody cloning, expression, and purification

Selected antibody sequences were purchased as gBlocks gene fragments (Integrated DNA Technologies) and cloned into a customized pcDNA3.4 vector (Invitrogen) containing human IgG1 or IgA1 Fc regions. VH and VL plasmids were transfected into 30 mL cultures of Expi293F cells (Invitrogen) at a 1:2 ratio and incubated at 37˚C and 8% CO_2_ for 7 days. The supernatant containing secreted antibodies was collected following centrifugation (1000 g for 10 min at 4˚C), neutralized, and filtered. Antibodies were isolated using Protein G Plus agarose (IgG; Pierce Thermo Fisher Scientific) or Peptide M agarose (IgA; Invitrogen) affinity chromatography, washed with 20 column volumes of PBS, eluted with 100 mM glycine-HCl pH 2.7, and immediately neutralized with 1 M Tris-HCl pH 8.0. The antibodies were then concentrated and buffer exchanged into PBS using 10,000 MWCO Vivaspin centrifugal spin columns (Sartorius).

#### Surface plasmon resonance

NY-ESO-1 (RayBiotech Inc.) was immobilized on CM5 sensor chips by amine coupling, as recommended by the manufacturer (GE Healthcare). Binding experiments were performed in HBS-EP buffer (10 mM HEPES pH 7.4, 150 mM NaCl, 3.4 mM EDTA, and 0.005% P20 surfactant).

Serially diluted antibodies were injected at a flow rate of 30 µL/min for 60 s with a dissociation time of 5 min. The chip was regenerated after each run by sequential injection of 10 mM glycine, pH 3, and 500 mM arginine, pH 8 for 1 min each. For each run, a bovine serum albumin (BSA)-coupled surface was used to subtract non-specific receptor binding. The K_D_ of each monoclonal antibody was calculated by fitting 2:1 bivalent analyte models (A+2B<->AB+B<->AB_2_) to the data using BIAevaluation 3.2 software (GE Healthcare) in accordance with previously reported analyses (42) and averaged from three independent experiments.

#### Recombinant expression of NY-ESO-1 for Ig-Seq

Full length NY-ESO-1 construct was provided by Dr. Richard Roden (Johns Hopkins University). This protein was expressed in a HIS-fusion bacterial expression system using the pET-28a expression vector (Novagen) as previously described (43). The cells from 1 L isopropyl-β-D-thiogalactopyranoside-induced bacterial cultures were lysed using 5 cycles of sonication followed by fusion protein solublization with 1% Tween-20 in PBS. The protein was purified under native conditions using HIS-Select(R) HF nickel affinity gel (Sigma) in batch processing format according to manufacturer’s instructions. The purity was >90%, as determined by SDS-PAGE gel treated with EZBlue Gel staining reagent (Sigma). Protein concentration was determined by Bradford-based protein assay (Biorad) with an average yield of 0.5 mg per liter.

## Acknowledgments

The authors thank the UVM Larner College of Medicine Microscopy Imaging Center for tissue imaging; the Genome Sequencing and Analysis Facility at the University of Texas at Austin for performing Illumina next-generation sequencing; the clinical coordinators in the Vermont Cancer Center for consenting patients and collecting the tissues for this study; Andrew D. Ellington (UT Austin) for a generous gift of RTX polymerase; and members of our laboratories for critical reading of the manuscript.

## Data Availability

The raw NGS data reported in this paper will be made available through the Sequence Read Archive (SRA). The raw proteomics data will be available through the mass spectrometry interactive virtual environment (MassIVE; http://massive.ucsd.edu).

## AUTHORS’ CONTRIBUTIONS

Conception and design: J.R. McDaniel, S.C. Pero, G.S. Shukla, N. Sidiropoulos, G.C. Ippolito, D. N. Krag Development of methodology: J.R. McDaniel, S.C. Pero, W.N. Voss, G.S. Shukla, J. Gollihar, J.W. Ellefson

Acquisition of data: J.R. McDaniel, S.C. Pero, W.N. Voss, Y. Sun, S. Schaetzle, C.H. Lee, C. C. Krag, Y.Tanno, S. Harlow

Acquired and Managed Patients: D.N. Krag and S. Harlow

Analysis and interpretation of data: J.R. McDaniel, S.C. Pero, W.N. Voss, G.S. Shukla, Y.Sun, S. Schaetzle,

C.H. Lee, A.P. Horton, G.C. Ippolito, D.N. Krag

Writing, review, and/or revision of the manuscript: J.R. McDaniel, S.C. Pero, W.N. Voss, G.S. Shukla, S. Schaetzle, C.H. Lee, A.P. Horton, G. Georgiou, G.C. Ippolito, D.N. Krag

Administrative, technical, or material support (i.e., reporting or organizing data, constructing databases):

S.C. Pero, W.N. Voss, J. Gollihar, J.W. Ellefson, Y. Tanno, G. Georgiou, D.N. Krag Study Supervision: J.R. McDaniel, S.C. Pero, G. Georgiou, G.C. Ippolito, D.N. Krag

## REFERENCES

1. Disis ML, Stanton SE. Immunotherapy in breast cancer: An introduction. The Breast [Internet]. 2017 Feb 3; Available from: http://www.sciencedirect.com/science/article/pii/S0960977617300139

2. Kroemer G, Senovilla L, Galluzzi L, André F, Zitvogel L. Natural and therapy-induced immunosurveillance in breast cancer. Nat Med. 2015 Oct;21(10):1128–38.

3. Adams S. Enlisting the Immune System to Cure Breast Cancer—A Recipe for Success. JAMA Oncol. 2016 Jan 1;2(1):25–7.

4. Nathan MR, Schmid P. The emerging world of breast cancer immunotherapy. Breast Edinb Scotl. 2017 Jun 2;

5. Krag DN. Minimal access surgery for staging regional lymph nodes: the sentinel-node concept. Curr Probl Surg. 1998 Nov;35(11):951–1016.

6. Krag DN, Anderson SJ, Julian TB, Brown AM, Harlow SP, Ashikaga T, et al. Technical outcomes of sentinel-lymph-node resection and conventional axillary-lymph-node dissection in patients with clinically node-negative breast cancer: results from the NSABP B-32 randomised phase III trial. Lancet Oncol. 2007 Oct;8(10):881–8.

7. Devarakonda CV, Kita D, Phoenix KN, Claffey KP. Patient-derived heavy chain antibody targets cell surface HSP90 on breast tumors. BMC Cancer. 2015 Sep 3;15:614.

8. Novinger LJ, Ashikaga T, Krag DN. Identification of tumor-binding scFv derived from clonally related B cells in tumor and lymph node of a patient with breast cancer. Cancer Immunol Immunother CII. 2015 Jan;64(1):29–39.

9. Williams LD, Ofek G, Schätzle S, McDaniel JR, Lu X, Nicely NI, et al. Potent and broad HIV-neutralizing antibodies in memory B cells and plasma. Sci Immunol. 2017 Jan 27;2(7):eaal2200.

10. Lavinder JJ, Horton AP, Georgiou G, Ippolito GC. Next-generation sequencing and protein mass spectrometry for the comprehensive analysis of human cellular and serum antibody repertoires. Curr Opin Chem Biol. 2015 Feb 1;24:112–20.

11. Wine Y, Horton AP, Ippolito GC, Georgiou G. Serology in the 21st century: the molecular-level analysis of the serum antibody repertoire. Curr Opin Immunol. 2015 Aug;35:89–97.

12. Chen YT, Scanlan MJ, Sahin U, Türeci O, Gure AO, Tsang S, et al. A testicular antigen aberrantly expressed in human cancers detected by autologous antibody screening. Proc Natl Acad Sci U S A. 1997 Mar 4;94(5):1914–8.

13. Cheever MA, Allison JP, Ferris AS, Finn OJ, Hastings BM, Hecht TT, et al. The prioritization of cancer antigens: a national cancer institute pilot project for the acceleration of translational research. Clin Cancer Res Off J Am Assoc Cancer Res. 2009 Sep 1;15(17):5323–37.

14. Hamaï A, Duperrier-Amouriaux K, Pignon P, Raimbaud I, Memeo L, Colarossi C, et al. Antibody responses to NY-ESO-1 in primary breast cancer identify a subtype target for immunotherapy. PloS One. 2011;6(6):e21129.

15. Theurillat J-P, Ingold F, Frei C, Zippelius A, Varga Z, Seifert B, et al. NY-ESO-1 protein expression in primary breast carcinoma and metastases: correlation with CD8+ T-cell and CD79a+ plasmacytic/B-cell infiltration. Int J Cancer. 2007 Jun 1;120(11):2411–7.

16. Yuan J, Adamow M, Ginsberg BA, Rasalan TS, Ritter E, Gallardo HF, et al. Integrated NY-ESO-1 antibody and CD8+ T-cell responses correlate with clinical benefit in advanced melanoma patients treated with ipilimumab. Proc Natl Acad Sci U S A. 2011 Oct 4;108(40):16723–8.

17. McDaniel JR, DeKosky BJ, Tanno H, Ellington AD, Georgiou G. Ultra-high-throughput sequencing of the immune receptor repertoire from millions of lymphocytes. Nat Protoc. 2016 Mar;11(3):429–42.

18. DeKosky BJ, Kojima T, Rodin A, Charab W, Ippolito GC, Ellington AD, et al. In-depth determination and analysis of the human paired heavy- and light-chain antibody repertoire. Nat Med. 2015 Jan;21(1):86–91.

19. Ellefson JW, Gollihar J, Shroff R, Shivram H, Iyer VR, Ellington AD. Synthetic evolutionary origin of a proofreading reverse transcriptase. Science. 2016 Jun 24;352(6293):1590–3.

20. Boutz DR, Horton AP, Wine Y, Lavinder JJ, Georgiou G, Marcotte EM. Proteomic identification of monoclonal antibodies from serum. Anal Chem. 2014 May 20;86(10):4758–66.

21. Lavinder JJ, Wine Y, Giesecke C, Ippolito GC, Horton AP, Lungu OI, et al. Identification and characterization of the constituent human serum antibodies elicited by vaccination. Proc Natl Acad Sci U S A. 2014 Feb 11;111(6):2259–64.

22. Wine Y, Boutz DR, Lavinder JJ, Miklos AE, Hughes RA, Hoi KH, et al. Molecular deconvolution of the monoclonal antibodies that comprise the polyclonal serum response. Proc Natl Acad Sci U S A. 2013 Feb 19;110(8):2993–8.

23. Rodriguez-Pinto D, Sparkowski J, Keough MP, Phoenix KN, Vumbaca F, Han DK, et al. Identification of novel tumor antigens with patient-derived immune-selected antibodies. Cancer Immunol Immunother CII. 2009 Feb;58(2):221–34.

24. Coronella JA, Spier C, Welch M, Trevor KT, Stopeck AT, Villar H, et al. Antigen-driven oligoclonal expansion of tumor-infiltrating B cells in infiltrating ductal carcinoma of the breast. J Immunol Baltim Md 1950. 2002 Aug 15;169(4):1829–36.

25. Spitzer MH, Carmi Y, Reticker-Flynn NE, Kwek SS, Madhireddy D, Martins MM, et al. Systemic Immunity Is Required for Effective Cancer Immunotherapy. Cell. 2017 Jan 26;168(3):487–502.e15.

26. Xu JL, Davis MM. Diversity in the CDR3 region of V(H) is sufficient for most antibody specificities. Immunity. 2000 Jul;13(1):37–45.

27. Ippolito GC, Schelonka RL, Zemlin M, Ivanov II, Kobayashi R, Zemlin C, et al. Forced usage of positively charged amino acids in immunoglobulin CDR-H3 impairs B cell development and antibody production. J Exp Med. 2006 Jun 12;203(6):1567–78.

28. Page DB, Yuan J, Redmond D, Wen YH, Durack JC, Emerson R, et al. Deep Sequencing of T-cell Receptor DNA as a Biomarker of Clonally Expanded TILs in Breast Cancer after Immunotherapy. Cancer Immunol Res. 2016 Oct 1;4(10):835–44.

29. Pasetto A, Alena G, Robbins PF, Deniger DC, Prickett TD, Matus-Nicodemos R, et al. Tumor- and neoantigen-reactive T-cell receptors can be identified based on their frequency in fresh tumor. Cancer Immunol Res. 2016 Jan 1;canimm.0001.2016.

30. Ademuyiwa FO, Bshara W, Attwood K, Morrison C, Edge SB, Karpf AR, et al. NY-ESO-1 cancer testis antigen demonstrates high immunogenicity in triple negative breast cancer. PloS One. 2012;7(6):e38783.

31. Rapoport AP, Stadtmauer EA, Binder-Scholl GK, Goloubeva O, Vogl DT, Lacey SF, et al. NY-ESO-1-specific TCR-engineered T cells mediate sustained antigen-specific antitumor effects in myeloma. Nat Med. 2015 Aug;21(8):914–21.

32. Robbins PF, Kassim SH, Tran TLN, Crystal JS, Morgan RA, Feldman SA, et al. A pilot trial using lymphocytes genetically engineered with an NY-ESO-1-reactive T cell receptor: Long term follow up and correlates with response. Clin Cancer Res Off J Am Assoc Cancer Res. 2015 Mar 1;21(5):1019–27.

33. Krag DN, Weaver DL, Alex JC, Fairbank JT. Surgical resection and radiolocalization of the sentinel lymph node in breast cancer using a gamma probe. Surg Oncol. 1993 Dec;2(6):335–339; discussion 340.

34. Krag DN, Anderson SJ, Julian TB, Brown AM, Harlow SP, Costantino JP, et al. Sentinel-lymph-node resection compared with conventional axillary-lymph-node dissection in clinically node-negative patients with breast cancer: overall survival findings from the NSABP B-32 randomised phase 3 trial. Lancet Oncol. 2010 Oct;11(10):927–33.

35. Bolger AM, Lohse M, Usadel B. Trimmomatic: a flexible trimmer for Illumina sequence data. Bioinformatics. 2014 Aug 1;30(15):2114–20.

36. Bolotin DA, Poslavsky S, Mitrophanov I, Shugay M, Mamedov IZ, Putintseva EV, et al. MiXCR: software for comprehensive adaptive immunity profiling. Nat Methods. 2015 May;12(5):380–1.

37. Plotly Technologies Inc. Collaborative data science. Montréal, QC, 2015. https://plot.ly.

38. Zhang J, Kobert K, Flouri T, Stamatakis A. PEAR: a fast and accurate Illumina Paired-End reAd mergeR. Bioinformatics. 2014 Mar 1;30(5):614–20.

39. Katoh K, Misawa K, Kuma K, Miyata T. MAFFT: a novel method for rapid multiple sequence alignment based on fast Fourier transform. Nucleic Acids Res. 2002 Jul 15;30(14):3059–66.

40. Stamatakis A. RAxML version 8: a tool for phylogenetic analysis and post-analysis of large phylogenies. Bioinformatics. 2014 May 1;30(9):1312–3.

41. Lee J, Boutz DR, Chromikova V, Joyce MG, Vollmers C, Leung K, et al. Molecular-level analysis of the serum antibody repertoire in young adults before and after seasonal influenza vaccination. Nat Med. 2016 Dec;22(12):1456–64.

42. Lee C-H, Romain G, Yan W, Watanabe M, Charab W, Todorova B, et al. IgG Fc domains that bind C1q but not effector Fcγ receptors delineate the importance of complement-mediated effector functions. Nat Immunol. 2017 Jun 12;

43. Tsai-Turton M, Santillan A, Lu D, Bristow RE, Chan KC, Shih I-M, et al. p53 autoantibodies, cytokine levels and ovarian carcinogenesis. Gynecol Oncol. 2009 Jul;114(1):12–7.

